# Glioblastoma Margin as a Diffusion Barrier Revealed by Photoactivation of Plasmonic Nanovesicles

**DOI:** 10.1101/2023.10.29.564569

**Authors:** Hejian Xiong, Blake A. Wilson, Xiaoqian Ge, Xiaofei Gao, Qi Cai, Xueqi Xu, Robert Bachoo, Zhenpeng Qin

## Abstract

Glioblastoma (GBM) is the most complex and lethal adult primary brain cancer. Adequate drug diffusion and penetration are essential for treating GBM, but how the spatial heterogeneity in GBM impacts drug diffusion and transport is poorly understood. Herein, we report a new method, photoactivation of plasmonic nanovesicles (PANO), to measure molecular diffusion in the extracellular space of GBM. By examining three genetically engineered GBM mouse models that recapitulate key clinical features including angiogenic core and diffuse infiltration, we found that the tumor margin has the lowest diffusion coefficient (highest tortuosity) compared with the tumor core and surrounding brain tissue. Analysis of the cellular composition shows that the tortuosity in the GBM is strongly correlated with neuronal loss and astrocyte activation. Our all-optical measurement reveals the heterogeneous GBM microenvironment and highlights the tumor margin as a diffusion barrier for drug transport in the brain, with implications for therapeutic delivery.

## Introduction

Glioblastoma (GBM) is the most common and aggressive malignant brain tumor among adults.^1^ The limited therapeutic delivery and the resistance to treatment have limited the progress in the field.^2^ Beyond the blood-brain barrier (BBB), the extracellular space (ECS) serves as a major obstacle for the drug to distribute in the tumor and diffuse to the targeted cells.^3^ Although some strategies could overcome or bypass the BBB, such as focused ultrasound^4^ or intracranial convection-enhanced delivery,^5^ effective drug accumulation in GBM is still challenging due to inefficient drug extravasation and penetration in tumors.^6, 7^ A major hallmark of GBM is the temporal and spatial heterogeneity, especially the cellular composition and extracellular matrix.^8^ Despite efforts to dissect the cellular and extracellular composition in GBM,^9, 10^ it remains poorly understood how molecules diffuse and transport in the heterogeneous GBM microenvironment.

So far, studies have shown the effect of extracellular matrix composition on the ECS diffusion in solid tumors, including GBM. For example, Sykova et al. measured the diffusion of tetramethylammonium (TMA^+^) ions in human gliomas using real-time iontophoresis and found that the tortuosity increases with tumor malignancy and extracellular glycoproteins deposition.^11, 12^ Jain et al. measured the diffusion of macromolecules in U87 glioblastoma xenograft by fluorescence recovery after photobleaching (FRAP) and found more hindered diffusion with high collagen type I content.^13^ Slowed diffusion and its correlation with matrix composition were also investigated using FRAP in several solid tumors, such as colon,^14^ lung^15^ and melanoma tumors.^13, 16^ Verkman et al. demonstrated that the slower diffusion of macromolecules deep in the tumor than near the surface in subcutaneously xenografted lung tumor.^15^ Although these studies have demonstrated diffusion could be an important barrier for drug delivery in tumors and it is well known that GBM is particularly heterogeneous, there have been very limited studies on the spatial heterogeneity of molecular diffusion in the tumor, despite its role in determining drug penetration and distribution.

This study focuses on addressing the heterogeneous transport properties in GBM in three preclinical GBM models that recapitulate key clinical features. Towards this, we present a novel technique utilizing the **P**hoto**A**ctivation of plasmo**N**ic nan**O**vesicles (PANO) to measure the molecular diffusion in the GBM extracellular space, which includes all-optical activation and observation of fluorophore diffusion. We applied the PANO technique to investigate the heterogeneous GBM brains in three preclinical genetically engineered glioma mouse models (GEMM), which carry mutations common in both adult and pediatric high-grade gliomas.^17, 18^ The orthotopic xenografted PS5A1 and 73C GBM were charactered with highly infiltrative margin and rapidly expanding core, respectively, which represent a reasonable facsimile of human GBM features. The third GEMM, de novo GBM, in a natural immune-proficient microenvironment more closely mimics the histopathological features and the progress of human GBM. In these three GEMMs, we found consistently that the tumor margin works as a barrier for molecular diffusion to the tumor core. Furthermore, analyzing the cellular composition reveals a strong correlation between the diffusion coefficient in GBM and neuronal loss as well as astrocyte activation. Our research offers fresh perspectives on the heterogeneous microenvironment of GBM and underscores the tumor margin as a diffusion obstacle for drug transport within the brain. This has significant implications for therapeutic delivery.

## Results

### Development and validation of the PANO technique for diffusion measurement in brain ECS

We first designed a workflow and validated the PANO technique in healthy brain tissue *in vitro* and *in vivo*. The workflow involves introducing the calcein-containing plasmonic nanovesicles into the samples, followed by short laser pulse excitation (720 nm), two-photon recording of the diffusion cloud (920 nm excitation), and image analysis to obtain the free diffusion coefficient (D), effective diffusion coefficient (D*) and tortuosity (λ, **Figure 1a**). Tortuosity is the measure of hindrance for molecule diffusion in ECS. We encapsulated calcein, a self-quenching polyanionic dye with minimal cellular uptake,^19^ in the plasmonic gold-coated nanovesicles (Au-nV-Cal). The gold coating absorbs energy from near-infrared laser pulses to trigger a rapid sub-second release.^20, 21^ Au-nV-Cal were characterized by UV-Vis spectrophotometry, dynamic light scattering, and transmission electronic microscopy (**Figure S1**). We then injected Au-nV-Cal (∼175 nm) into 0.2% agarose gel as a “free” medium (**Figure 1b**).^22, 23^ Since calcein fluorescence is self-quenched in the nanovesicles due to the high concentration (75 mM), the photoreleased calcein (∼70 µM) shows a bright fluorescence increase and could be considered as the point source for diffusion measurement (**Figure 1b**). 2D Gaussian function fitting (Equation 1, the lower panel of **Figure 1b**) captures the diffusion cloud changes over time. Further linear fitting (Equation 2) gives the free diffusion coefficient (D=42.9±2.5 × 10^-7^ cm^2^•s^-1^) for calcein from this measurement at 31 °C (**Figure 1c**).

**Figure 1.**
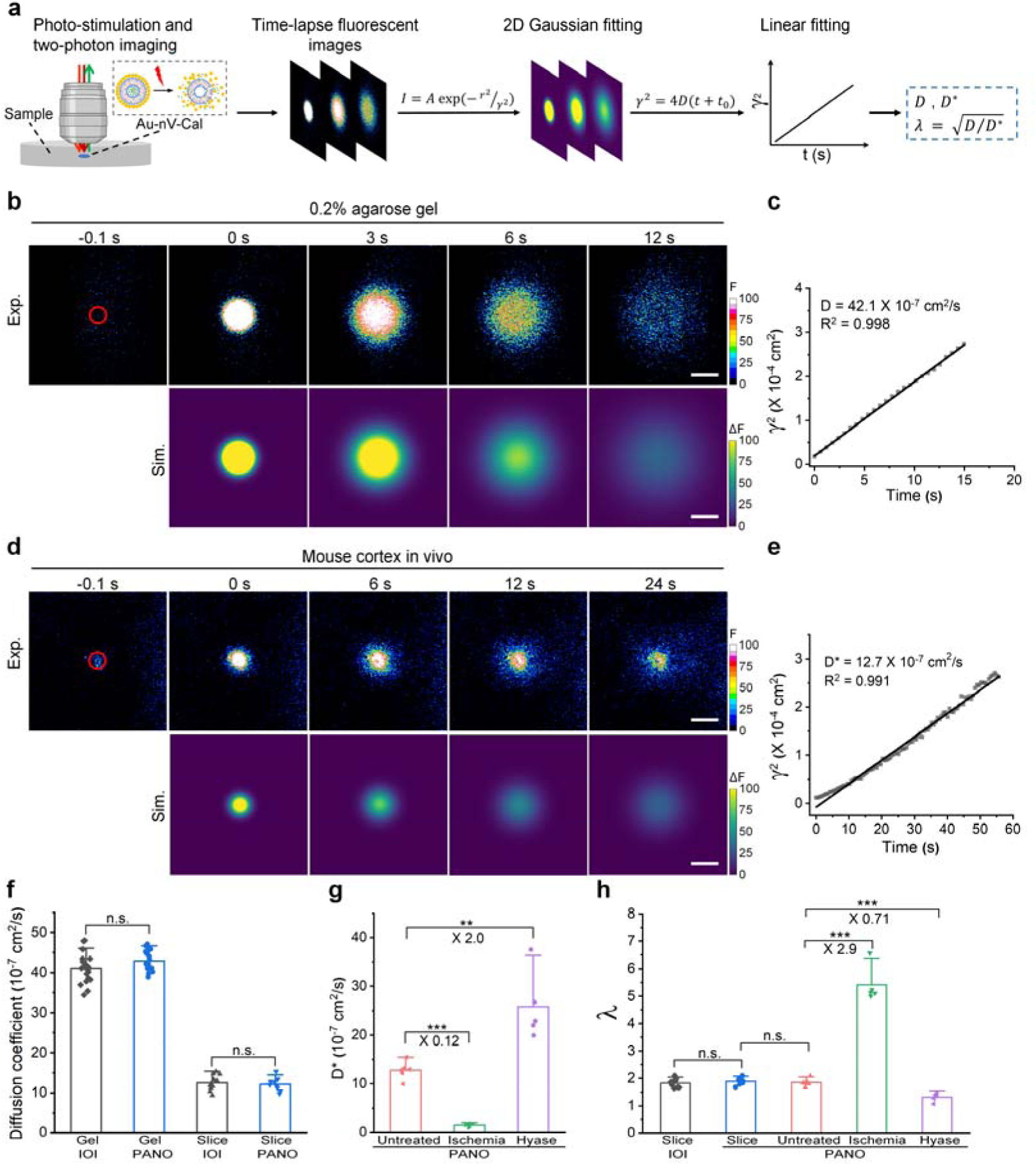
Workflow and characterization of the PANO technique *in vitro* and *in vivo*. (a) Experimental and analysis workflow of the PANO technique. Au-nV-Cal was injected into the samples and photo-stimulated by 720 nm laser pulses. Time-lapse two-photon imaging of released calcein was synchronized after stimulation and fitted by a 2D Gaussian equation. The final linear fitting gives the diffusion parameters. I, r, and t are the fluorescence intensity, distance to the point source, and time, respectively. A and γ are determined by the fitted images to the fluorescent images with the Nelder-Mead algorithm. (b) Two-photon fluorescent images (upper panel) and 2D Gaussian fitted images (lower panel) of calcein diffusion in 0.2% agarose gel. 60-µm-diameter tornado scans (red circle) were performed on Au-nV-Cal for 0.1 s before the diffusion recording. Scale bar: 100 µm. (c) Linear fitting of γ^2^ versus *t* to get the diffusion coefficient. (d) Two-photon fluorescent images (upper panel) and 2D Gaussian fitted images (lower panel) of calcein diffusion in the mouse cortex *in vivo*. 60-µm-diameter tornado scans (red circle) were performed on Au-nV-Cal for 0.1 s before the diffusion recording. Scale bar: 100 µm. (e) Linear fitting of γ versus *t* to get the diffusion coefficient. (f) Comparison of calcein diffusion coefficients in 0.2% agarose gel and acute brain slice measured by PANO and integrative optical imaging (IOI) methods. IOI: n=24 recordings in agarose gel and n=11 slices of cortex from 5 mice. PANO: n=18 recordings in agarose gel and n=10 slices of cortex from 3 mice. (g) Calcein diffusion coefficients *in vivo* measured by PANO in different mouse models. Untreated: n=6 independent implants in the cortex from 3 mice; Ischemia: n=5 independent implants in the cortex from 5 mice; Hyase: n=5 independent implants from 3 hyaluronidase treated mice. (h) Tortuosity (λ) of calcein in the diluted agarose gel and cortex at different conditions compared with the results from the IOI method. Statistical analysis was performed by two-sample Student’s t-tests. Data are expressed as Mean ± S.D.; \*\**p* < 0.01; \*\*\**p* < 0.001; n.s., not significantly different.

To validate the PANO method *in vivo*, we then injected Au-nV-Cal into the mouse somatosensory cortex, which was imaged within 2-4 hours at 200 µm below the surface (cortical layer II) through an open cranial window. Upon 720 nm femtosecond laser stimulation, representative two-photon image sequences clearly show that extracellular diffusion of calcein dye is more hindered in the mouse cortex than in the agarose gel (**Figure 1d** vs **Figure 1b**). Data analysis gives an effective diffusion coefficient for calcein as D*=12.8±1.8 × 10^-7^ cm^2^•s^-1^ in the mouse cortex at 31 °C (**Figure 1e**). The measured calcein diffusion coefficients in 0.2% agarose gel and cortex of brain slices obtained with the PANO method are benchmarked and consistent with the data obtained with the well-established integrative optical imaging (IOI) method (**Figure 1f**). The tortuosity value (1.8 ± 0.1) agrees well with the IOI measurement in the cortex of brain slices (**Figure 1h**) and previous literature.^24^ Further testing on brain ischemia under cardiac arrest shows that D* decreased to 12% of its normoxic value, while the tortuosity (λ) was 2.9-fold higher (**Figure 1g and 1h, Figure S2**). On the other hand, the degradation of hyaluronan, a major extracellular matrix component, leads to 2.0-fold increase in the D* and 71% decrease in tortuosity compared with untreated brains (**Figure 1g and 1h**). These results agree with the report that cardiac arrest could cause the swelling of brain cells and the shrinkage of ECS to decrease extracellular diffusion,^25–27^ while hyaluronan degradation increases the effective diffusion coefficient.^28, 29^ The results confirm the feasibility, accuracy, and versatility of our all-optical PANO method for diffusion measurement.

### Spatially heterogeneous molecular diffusion in three GBM models

Next, we examined the heterogeneous diffusion properties of GBM. GBM is a highly heterogeneous disease characterized by a solid, often necrotic core and an infiltrative margin.^2, 8^ We first tested two genetically engineered mouse models (GEMMs) of glioma by transplanting primary conditional mouse astrocyte and neural stem cell lines into mouse brains (**Figure 2a**). The first model uses PS5A1 cell line contained common Braf^V600E^, INK4ab/Arf^-/-^, PTEN^-/-^ mutations in both adult and pediatric high-grade gliomas with tdTomato markers.^17^ PS5A1 GBM is characterized by the gradual decrease of glioma cell density towards the brain parenchyma due to the high infiltration (**Figure 2b**). The glioma margin is defined as the region within 500 µm from the tumor core (glioma cell density > 3/500 µm^2^) in this work. We injected the Au-Cal-nV into the acute brain slices of PS5A1 GBM. Interestingly, we found that the calcein diffusion in the tumor margin is 72% slower compared with the cortex, while the diffusion coefficient in the tumor core is 1.6-fold higher than that in the tumor margin (**Figure 2c**). The tortuosity in the tumor margin is the highest compared with the cortex and tumor core (**Figure 2d**), indicating that the tumor margin is the main barrier for molecular diffusion.

**Figure 2.**
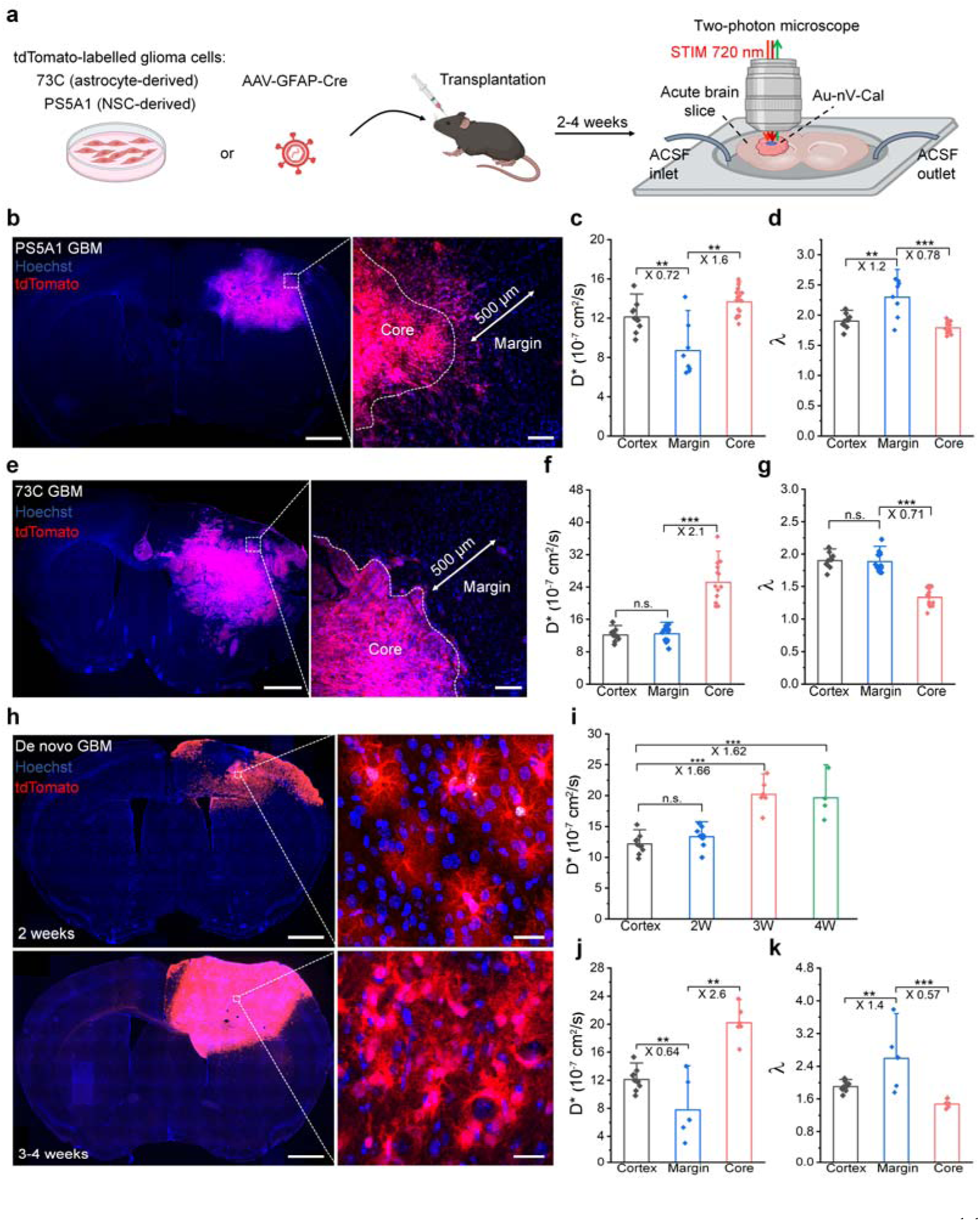
Spatially-resolved diffusion measurement by the PANO technique in three orthotopic GBM models. (a) Schematic of establishing the orthotopic 73C, PS5A1 and de novo GBM mouse models and the diffusion measurement in the acute brain slices. NSC: neural stem cell. (b) Fluorescent images of PS5A1 GBM. Scale bar: 1 mm (left) and 50 μm (right). (c) Diffusion coefficient and (d) tortuosity of calcein dye in the PS5A1 GBM. (e) Fluorescent images of 73C GBM and definition of the tumor core and margin. Scale bar: 1 mm (left) and 50 μm (right). (f) Diffusion coefficient and (g) tortuosity of calcein in the 73C GBM including core, margin, and contralateral cortex. (h) Progression of de novo GBM. Scale bar: 1 mm (left) and 20 μm (right). (i) Calcein diffusion coefficient during the progression of de novo GBM. (j) Diffusion coefficient and (k) tortuosity of calcein in the de novo GBM. Red: tdTomato-labelled glioma cells; blue: Hoechst 33342-labeled cell nuclei. Data are expressed as Mean ± S.D; \*\**p* < 0.01; \*\*\**p* < 0.001; n.s., not significantly different.

We then tested 73C GBM that carries mutations including Braf^V600E^, P53^-/-^, and PTEN^-/-^, seen in both adult and pediatric high-grade gliomas and is derived from conditional multi-allele primary astrocytes.^17, 18^ 73C GBM is characterized by a clear boundary between the tumor core and margin (**Figure 2e**), resulting from a fast-growing angiogenic tumor with limited infiltration. We found that the diffusion coefficient in the tumor core is 2.1-fold higher than that in the tumor margin and the tortuosity of ECS in the tumor core is around 71% of that in the tumor margin (**Figure 2f,g, Figure S3**). This demonstrates that molecules can easily diffuse into the tumor core, while face larger resistance towards the tumor margin.

We further applied the technique to investigate the ECS of a de novo GBM model. There has been significant interest in de novo models of GBM in which the tumor evolves in a natural immune-competent microenvironment reflecting the crosstalk of cancer cells with the tumor microenvironment (including infiltrating immune cells, fibroblasts, and the glymphatic and blood vasculature) as observed in human cancer.^30, 31^ The de novo GEMMs can closely mimic the tumor progression and display genetic heterogeneity, as well as the histopathological and molecular features of their human counterparts.^32^ For this model, AAV-GFAP-Cre was injected into the transgenic mice to induce the mutation and malignant proliferation of instinct astrocytes (**Figure 2a**). We monitored the morphology changes of astrocytes and measured the ECS diffusion during the tumor progression. The astrocytes are labeled by tdTomato in the injection area. The number and length of astrocytic processes decrease during tumor growth (**Figure 2h**). The diffusion coefficient of calcein in the tumor core increases after two weeks of initiation and remains constant in 3- and 4-week GBM (**Figure 2i**). The diffusion coefficient in the tumor margin is 64% of that in the cortex and 38% of that in the tumor core (**Figure 2j**), and the tortuosity in the tumor margin is highest (**Figure 2k**).

### Cellular heterogeneity in GBM and correlation analysis

To better understand the cellular composition in different GBM regions, we labeled neurons, astrocytes, and microglia in the brain by immunostaining. **Figure 3a** shows that neurons are largely absent in the PS5A1 core. In contrast, a large amount of microglia appears in the tumor core, and there are abundant astrocytes and microglia in the PS5A1 margin. Quantitative analysis shows that the number of neurons in the tumor margin is close to that in the contralateral side in PS5A1 GBM (**Figure 3b**), while the number of functional synapses in the tumor margin is only 36% of that in the contralateral side (**Figure S4**). The number of astrocytes and microglia in the tumor core is around 4-fold and 110-fold higher than the contralateral side in PS5A1 GBM (**Figure 3b**). For 73C GBM, we observed minimal neurons and astrocytes in the tumor core and significantly increased astrocytes and microglia in the tumor margin (**Figure 3c,d**, **Figure S5**). Similar to PS5A1 GBM, there is largely an absence of neurons in the de novo GBM core. In contrast, the number of astrocytes and microglia in the tumor core is both over 100-fold higher than the contralateral side (**Figure 3e,f**, **Figure S5**). The enrichment of astrocytes in the tumor core could also be contributed by the immunostaining of malignant proliferative astrocytes. The number of reactive astrocytes and microglia in the de novo GBM margin is also 12-fold and 26-fold higher than in the contralateral cortex (**Figure 3e,f**).

**Figure 3.**
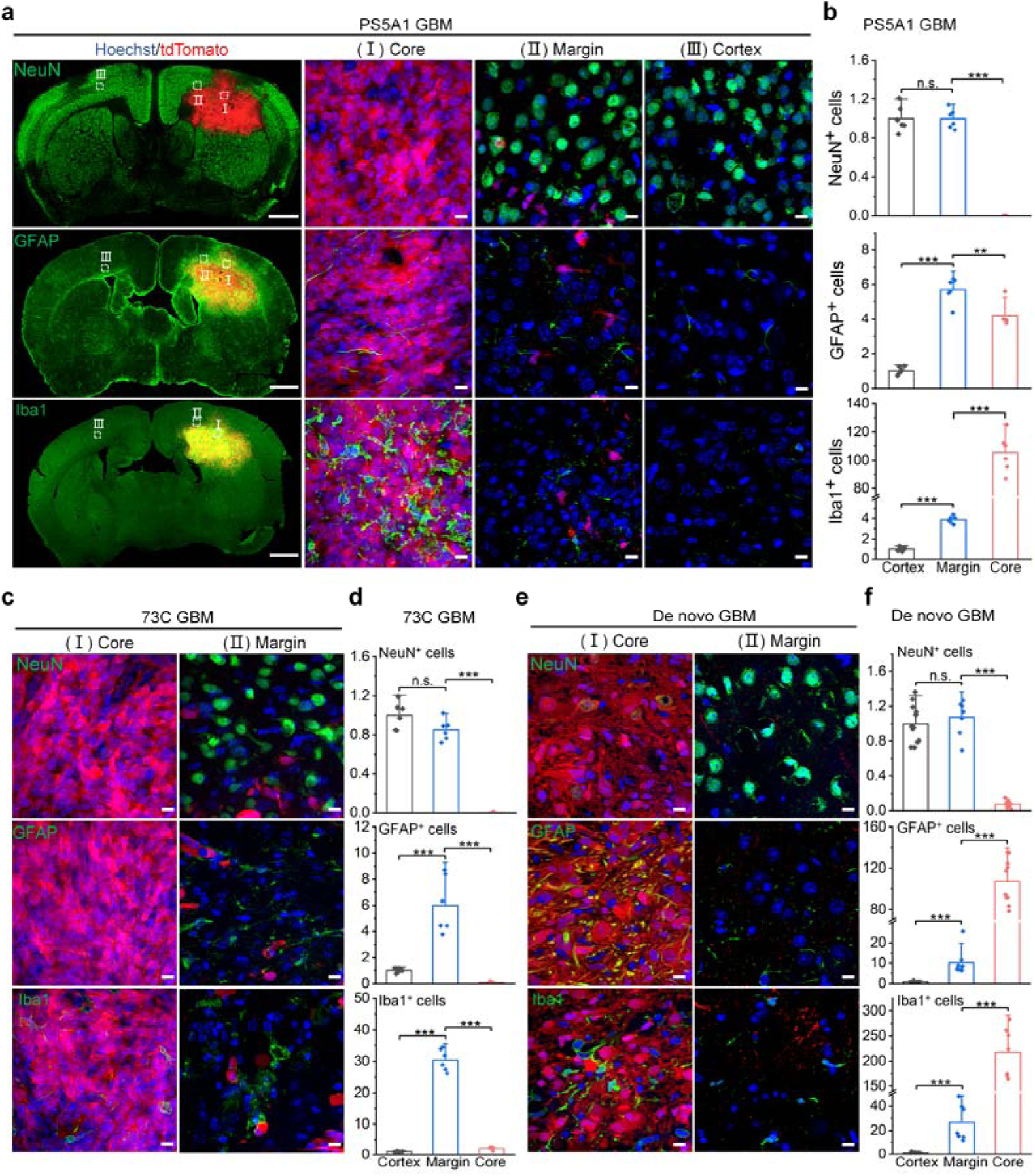
Characterization of cellular compositions in the GBMs. (a) Images of immunostained brain cells in PS5A1 GBM. Scale bar: 1 mm (left column) and 10 µm (right three columns). (b) Quantitative analysis of the immunostained brain cells in different regions of 3-week PS5A1 GBM (n=3 mice). The fluorescent area of each stained cell in the field of view (133 µm × 133 µm) was normalized to that of the contralateral cortex. (c) Images of immunostained brain cells in different regions of 3-week 73C GBM. Nuclei were labeled by Hoechst 33342 (blue). Neurons, astrocytes, and microglia were labeled by Alexa Fluor 488 (green). Scale bar: 10 µm. (d) Quantitative analysis of the immunostained brain cells in different regions of 3-week 73C GBM (n=3 mice). The fluorescent area of each stained cell in the field of view (133 µm × 133 µm) was normalized to that of the contralateral cortex. (e) Images of immunostained brain cells in different regions of 3-week de novo GBM. Scale bar: 10 μm. (f) Quantitative signal analysis from immunostained brain cells in 3-week de novo GBM. The fluorescent area of each stained cell in the field of view (133 µm × 133 µm) was normalized to that of the contralateral cortex. In figure a,c,e, nuclei were labeled by Hoechst 33342 (blue); neurons, astrocytes, and microglia were labeled by Alexa Fluor 488 (green); glioma cells were labeled by tdTomato (red). Statistical tests were performed by two-sample Student’s t-tests. Data are expressed as Mean ± S.D.; \*\**p* < 0.01; \*\*\**p* < 0.001; n.s., not significantly different.

Lastly, we performed a correlation analysis of our dataset on these three GBM tumor models. We summarized all the data, including diffusion coefficients and cell counting from the GBM in **Table S1**. We calculated the correlation coefficient using the Spearman rank correlation as a measure of a monotonic association, which gives a dimensionless measure of the covariance ranging from −1 to +1.^33^ **Table 1** shows that there is a strong positive correlation between the tortuosity and the density of neurons and astrocytes. however, the tortuosity has a low correlation with the microglia density.

**Table 1.** Spearman’s rank correlation analysis between tortuosity λ and cell density.

|  | Neurons | Astrocytes | Microglia |
| --- | --- | --- | --- |
| $\lambda$ | 0.86 | 0.71 | -0.25 |

## Discussion

Our study introduces the PANO method to measure molecular diffusion in the brain using photoactivated nanovesicles and 2-photon microscopy. With this tool, we discovered the diffusion barrier for the first time in the tumor margin between the healthy brain tissue and the tumor core in three preclinical genetically engineered GBM mouse models. These GBM recapitulate GBM features and progression, including the diffusely infiltrative tumor margin and angiogenic core. Furthermore, the diffusion properties in these GBM are strongly correlated with neuronal loss and astrocyte activation. Our study has several implications for the field as discussed below.

First, the PANO method opens new avenues for diffusion measurement in the brain ECS. The method enables the optical scanning of the plasmonic nanovesicles and creating point sources for diffusion measurement with good temporal and spatial resolution. Diffusion-weighted MRI can assess the glioma malignancy and treatment response by generating apparent diffusion coefficient (ADC) maps in the clinic.^34, 35^ However, it averages the water diffusion in both the intracellular and extracellular spaces, which prevents the quantitative analysis of the ECS properties in GBM.^24, 36^ While fluorescence recovery after photobleaching (FRAP) is the closest method to create photobleaching and measure molecular diffusion in tumors,^37^ its implementation for in vivo measurement has been limited since the fluorophore is constantly diffusing and diluted. On the contrary, the plasmonic nanovesicles are immobilized upon injection into the brain due to their large size.^22^ The PANO method also offers easier control to introduce fluorescent probes into the samples by a remote laser pulse trigger.

Second, our study reveals the GBM margin as a significant diffusion barrier. Previous studies have shown correlations between ECS diffusion, tumor type, and extracellular matrix compositions, while the spatial heterogeneity of ECS diffusion in solid tumors, especially in GBM, is largely unknown. Here, our study in the three GEMMs demonstrates that the diffusion is slowest in the tumor margin compared with tumor core and the surrounding healthy tissue, suggesting the tumor margin as diffusion barrier (**Figure 4**). We further examined the extracellular matrix and observed overexpression of collagen IV, Tenascin-C, and fibronectin in the core of 73C GBM, while the hyaluronan level in GBM is similar compared with other brain regions (**Figure S6**). The extracellular matrix deposition does not correlate with an increase in diffusion in the tumor core. On the other hand, the increase in extracellular volume fraction can enhance diffusion^38^. The fast diffusion in the tumor core might be due to the increase of ECS volume associated with neuronal loss (**Figure 3**), while the slow diffusion in the GBM margin may be contributed by the decrease of ECS volume with astrocyte activation. The discovery of the tumor margin as a diffusion barrier is valuable to better understand the tumor heterogeneity and devise strategies for effective therapeutic delivery.

**Figure 4.**
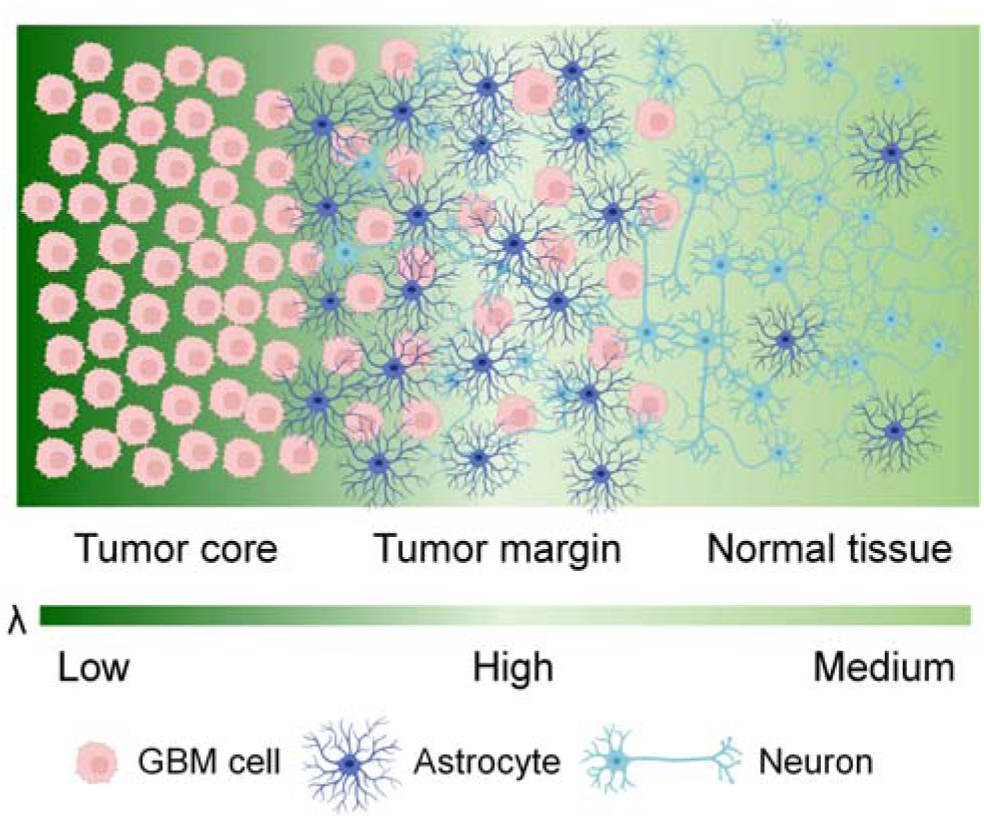
Summary of the heterogeneous GBM microenvironment in the brain. Tumor margin represents a diffusion barrier with high tortuosity (λ).

Third, we have evaluated preclinical animal GBM models that recapitulate the complexity of human GBM. GBM is highly heterogeneous with angiogenic core and infiltrative margin.^39^ To capture the features, we first tested two genetically engineered GBM cell lines (PS5A1 and 73C) to establish GEMMs. The GEMMs contain the common mutations in human GBM, e.g., the loss of critical tumor suppressor genes (PTEN^-/-^ and P53^-/-^; or PTEN^-/-^ and INK4ab/Arf^-/-^). We have characterized the heterogeneous vascular properties of the two GEMMs in the previous work.^18^ Here, we demonstrate the spatially cellular heterogenicity in these GEMMs, including the loss of neurons in the tumor core and the activation of astrocytes in the tumor margin. Moreover, the de novo GEMM includes the mutations Braf^V600Ef/+^; P53^f/f^; Pten^f/f^ seen in both adult and pediatric high-grade GBM. The de novo GBM could spontaneously progress in a natural immune-proficient microenvironment and closely mimic the histopathological and molecular features of human GBM.^40^ Taken together, these GEMMs are more relevant preclinical models to investigate the transport of the molecules in the tumor.

## Conclusion

In summary, we developed an all-optical PANO approach to measure the molecular diffusion in the brain and demonstrated the spatially heterogeneous ECS diffusion in GBM. We validated that the PANO technique is robust in acute brain slices and in vivo with different mouse models. Using the technique, we found that the tumor margin acts as a diffusion barrier between the healthy brain tissue and the tumor core in three preclinical genetically engineered GBM mouse models. We further demonstrated that the spatially resolved diffusion property is strongly correlated with neuron loss in the tumor core and astrocyte activation in the margin. Therefore, the PANO technique is useful in investigating the transport properties in the heterogeneous GBM microenvironment, and our findings provide novel insight with implications for improved therapeutic delivery.

## ASSOCIATED CONTENT

### Supporting Information

The following files are available free of charge.

Additional experimental materials and methods and additional figures, including glioma cell culture, preparation of open-skull cranial window, glioma cell transplantation, establishment of de novo GBM model, brain slice preparation, Au-nV implantation, two-photon imaging and stimulation, immunohistochemistry staining, analysis of diffusion coefficient, statistical analysis; and additional figures and table, including characterization of Au-nV-Cal, Calcein diffusion measurement in the somatosensory cortex of ischemic and hyase-treated mouse brains, Calcein diffusion in 73C GBM, immunostaining of synapses, images of immunostained brain cells in the whole slice of 73C and de novo GBMs, differential expression of extracellular Matrix (ECM) proteins in the 73C GBM, summary of cellular heterogeneity and tortuosity (PDF)

## AUTHOR INFORMATION

### Author Contributions

H.X., R.B. and Z.Q. conceived and designed the experiments. B.A.W. developed the code for diffusion analysis and the theoretical modeling for molecular diffusion in brain ECS. X. Ge performed the ECS imaging by Spinning Disk microscopy and assisted in the images analysis. X. Gao established the PS5A1 cell lines and the de novo glioma model. X.X. and Q.C. assisted in glioma cells transplantation and immunohistochemistry staining. H.X. and X. Ge performed the diffusion measurements and immunohistochemistry staining, analyzed the results and wrote the manuscript. H.X., B.A.W., X. Ge, R.B. and Z.Q. discussed the results and revised the manuscript. R.B. and Z.Q. supervised the project. All authors discussed the results and commented on the manuscript.

### Notes

The authors declare no competing financial interests.

## ACKNOWLEDGMENT

We thank Prof. Sabina Hrabetova from SUNY Downstate Health Sciences University for the calcein diffusion measurement by integrated optical imaging method and helpful comments on the manuscript. This work was partially supported by National Science Foundation under award number 2123971 (Z.Q.), National Institute of Neurological Disorders and Stroke of the National Institutes of Health (NIH) under award number RF1NS110499 (Z.Q.), Cancer Prevention, and Research Institute of Texas (CPRIT) grants RP190278 (Z.Q.). Some schematics were made in part using BioRender.

---

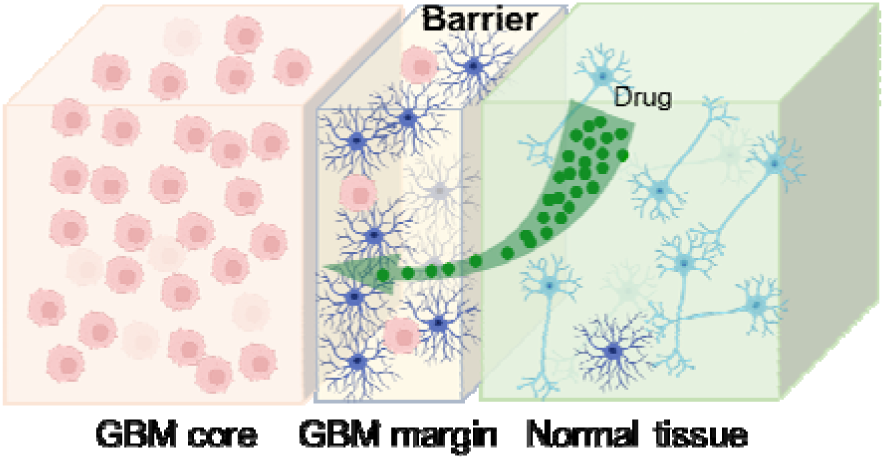
For Table of Contents only.

