## Supplemental methods and figures for "Glioblastoma Margin as a Diffusion Barrier Revealed by Photoactivation of Plasmonic Nanovesicles"

**Materials**:

1,2-Dipalmitoyl-*sn*-glycero-3-phosphocholine (DPPC, 734 g/mol, 63-89-8, >99%) and cholesterol (386.6 g/mol, 57-88-5, >98%) were purchased from Avanti Polar Lipids, Inc. Calcein sodium salt (644.5 g/mol, 108750-13-6) was purchased from Honeywell Fluka. Tetrachloroauric(III) trihydrate (HAuCl_4_·3H_2_O, 16961-25-4, 99.9%) and *L*-ascorbic acid (50-81-7, 99%), epidermal growth factor (EGF, Cat. No. E4127), fibroblast growth factor-basic human (FGF2, Cat. No. F0291), progesterone (Cat. No. P6149) were purchased from Sigma-Aldrich. Gibco™ Dulbecco’s Modified Eagle’s Medium (DMEM, high glucose, Cat. No.10569-010) with GlutaMAX™ supplement, DMEM/Hams F12 50:50 Mix (DMEM/F12, Cat. No. 15-090-CV), trypsin-EDTA solution (0.25%), fetal bovine serum (FBS), StemPro™ Accutase™ cell dissociation reagent, B-27TM supplement, insulin transferrin solution and penicillin-streptomycin were purchased from ThermoFisher Scientific. Gold-coated calcein-loaded liposomes (Au-nV-Cal) was prepared according to previous work.^1^ The hydrodynamic size and morphology of Au-nV-Cal was determined by dynamic light scattering measurement (Malvern Zetasizer Nano ZS) and a transmission electron microscope (TEM, JEOL-1400+) at an accelerating voltage of 150 keV, respectively. All other chemicals were analytical grade.

**Glioma cell culture**

73C and PS5A1 glioma cells were obtained from Dr. Robert Bachoo’s laboratory in the University of Texas Southwestern Medical Center. 73C glioma cells were generated in primary astrocyte cultures from neonatal mice that carried conditional mutations for Pten^f/f^, p53^f/f^, and LSLBraf^V600E^.^2^ 73C glioma cells were cultured in DMEM with 10% FBS and 1% penicillin-streptomycin. PS5A1 is a highly invasion tumor phenotype derived from de novo glioma in the adult BL6 background conditional mouse (Braf^V600Ef/+^; Ink4ab/Arf^f/f^; Pten^f/f^) that was induced by intracranial injection of AAV5-GFAP-Cre-tdTomato. The primary neurosphere cultures were then infected with a Lenti-tdTomato and selected by puromycin with stable red fluorescent protein expression. PS5A1 neurospheres were cultured in DMEM/F12 medium with 2% B-27, 1% insulin transferrin solution, 20 ng/mL of EGF, 20 ng/mL of FGF2, and 20 ng/mL progesterone. Glioma cells were cultured in an incubator at 37 ^o^C with 5% CO_2_.

**Animals**

7-week-old C57BL/6 mice were purchased from Charles River Laboratories, Inc., and 6-week-old homozygous nude mice *Foxn1^nu^* (Strain No. 002019) were purchased from the Jackson Laboratory. All mice were given free access to food and water before the experiments. Animal protocols were in accordance with NIH guidelines and approved by the Institutional Animal Care and Use Committee of the University of Texas at Dallas, SUNY Downstate Health Sciences University, and the University of Texas Southwestern Medical Center. For hyaluronan-deficient mouse model, approximately 4 μL of 20 mg/mL hyaluronidase from bovine testes in PBS (#H3506, Sigma Aldrich) was injected intraventricularly at the coordinate of -0.5 mm A/P, +1.0 mm M/L and a depth of 2 mm two days before the diffusion measurement. For terminal ischemia, mice were administered 1 mL of 1M KCl intracardially to induce immediate cardiac arrest.

**Preparation of open-skull cranial window**

C57BL/6 mice were anesthetized with 1.5-2% isoflurane (Vedco, Inc) and fixed in a stereotaxic frame with ear bars. A square piece of the skull (~3×3 mm) above the somatosensory cortex was removed to prepare an open-skull cranial window for microinjection and imaging. A square groove was first thinned slowly with a drill (EXL-M40, OSADA), followed by the lift of the central island of the skull bone. The implantation of Au-nV-Cal was conducted immediately after the surgery. The mouse body temperature was maintained at 37 °C using a heating pad regulated by a rectal probe. Dexamethasone (Sigma-Aldrich) was administered subcutaneously at 5 mg/kg to prevent swelling of the brain and/or an inflammatory response. Buprenorphine SR-Lab (ZooPharm) was administered subcutaneously at 1 mg/kg for postoperative analgesia.

**Glioma cell transplantation**

Nude mice *Foxn1^nu^* were anesthetized with 1.5-2% isoflurane and fixed in a stereotaxic frame with ear bars. A small window with a diameter of around 0.5 mm was created on the skull with a drill at the following stereotaxic coordinates: 0 mm A/P, -2 mm M/L, relative to Bregma. Glioma cells (73C and PS5A1) were harvested when reaching 70-80% confluence. Cells were dissociated, resuspended in Hank’s Balanced Salt Solution (HBSS, without Ca^2+^ or Mg^2+^), and filtered through a Cell Strainer (Corning^TM^) with a pore size of 70 µm. After counting, a cell suspension was prepared at a density of 2×10^5^/µL and loaded into a glass micropipette with a 50 µm tip generated by a Micropipette Puller (P-1000, Sutter Instrument Co.). Around 180 nL of glioma cells were injected into the mouse cortex at a depth of 1 mm using a Nanoliter2010 injector (World Precision Instruments). Buprenorphine SR-Lab was administered subcutaneously at 1 mg/kg for postoperative analgesia. The mice were housed for 2-3 weeks to allow glioma growth.

**Establishment of de novo GBM model**

De novo mouse glioma, in this specific test, was induced in adult mixed background conditional mouse with multiple floxed genes (Braf^V600Ef/+^; P53^f/f^; Pten^f/f^), which were frequently mutated or deleted in human glioma cases, with tdTomato reporter gene, by intracranial injection of GFAP promoter driven Adeno Associated Virus with Cre recombinase DNA in the cerebral cortex. Viral injection was carried out by using 1 µL Hamilton syringe with replaceable 34-gauge needle. With 250 nL of AAV5-GFAP-Cre (Addgene #105550-AAV5, titer ≥ 7×10¹² vg/mL) loading in the syringe that was mounted on motorated mini pump. AAV was gently delivered at the speed of 100 nL/min to the targeting location (coordinate: -2mm, 0.5mm, -1.5mm) of the mouse brain. Infected GFAP^+^ cell populations were transformed into highly proliferative and malignant tumor mass during the latency of 5-6 weeks after viral injection.

**Brain slice preparation**

Acute brain slices from glioma-bearing mice were prepared to measure the calcein diffusion. Mice were anesthetized with 5% isoflurane and then decapitated to extract the brain. The brain was submerged in artificial cerebrospinal fluid (ACSF, containing 125 mM NaCl, 2.5 mM KCl, 26 mM NaHCO_3_, 1.25 mM NaH_2_PO_4_, 10 mM D-Glucose, 1.5 mM MgCl_2_ and 2 mM CaCl_2_, pH 7.3, continuously bubbling with a mixture of 95% O_2_ and 5% CO_2_) precooled to 4 °C. Coronal slices with a thickness of 400 μm were prepared using a vibratome (Leica VT1200S) and kept in ACSF at 34 °C for 30 min. The slices were then allowed to recover at room temperature for 30 min. Afterwards, the slices were transferred onto a slice chamber (Cat. No. RC-26G, Warner Instrument) and anchored by a grid (Cat. No. 640255, Warner Instrument) for imaging. The slice was perfused with ACSF at a rate of 0.5 mL/min and kept at 31 ± 1 °C by a temperature controller (TC-344C, Warner Instrument). ACSF was continuously bubbled with 95% O_2_ and 5% CO_2_ during the whole process.

**Au-nV-Cal implantation**

Nanoliter2020 injector (World Precision Instruments) was used to conduct all the microinjections of Au-nV-Cal into the samples. Au-nV-Cal was loaded into a glass micropipette with a tip diameter of 30 µm. The injection volume was controlled by the internal micrometer step motor in the injector at a speed of 1 nL/s. To measure the free diffusion coefficient (*D*), the photorelease and monitoring of calcein were performed in 0.2% agarose gel prepared with 0.01 M PBS, which could be considered as a ‘‘free’’ medium.^3, 4^ Au-nV-Cal was loaded into a glass micropipette with a tip diameter of 30 µm, which was connected to a Nanoliter2020 injector (World Precision Instruments). The injection volume was controlled by the internal micrometer step motor in the injector at a speed of 1 nL/s. Finally, around 20 nL of Au-nV-Cal was injected into the gel. After implantation, the agarose gel was kept at 30±0.5 ^o^C by an incubator (Tokai Hit).

For the *in vivo* diffusion measurement, around 30 nL of Au-nV-Cal was injected into the somatosensory cortex 400-500 µm below the cortical surface through the open-skull cranial window within the following stereotaxic coordinates: -1.5 to 1.5 mm A/P, +1.5 to +2.5 mm M/L, relative to Bregma. After implantation, the mice were immobilized on a custom-built imaging frame and taken for two-photon imaging immediately.

For the diffusion measurement in acute brain slices, around 20 nL of Au-nV-Cal was injected into target areas, such as the core or the margin of the GBM at a depth of 200-300 µm. After implantation, photo-stimulation and imaging were performed immediately.

**Two-photon imaging and stimulation**

A multi-photon laser scanning microscope (FVMPE-RS, Olympus) accompanied with a stimulation laser (MaiTai HP DeepSee-OL, Spectra-Physics, 100 fs pulse width) and a imaging laser (Insight DS+ -OL, Spectra-Physics, 120 fs pulse width) with a repetition rate of 80 MHz was used for stimulation and imaging of Au-nV-Cal. Calcein fluorescence emission within 495-540 nm was excited at 920 nm and recorded by a 25X objective (XLPLN25XWMP2, Olympus, NA 1.05) at 0.6 s per frame. Photo-stimulation on the nanovesicles was performed by 5 tornado scans (diameter: 30 µm; duration 0.1 s) at 720 nm (75 or 100 mW). Here tornado scan refers to a scan setting for fast photo-stimulation and allows rapid scanning of a circular area. The focus plane was at 150-200 μm below the surface of samples.

**Immunohistochemistry staining**

To stain neurons (NeuN), astrocytes (GFAP), and microglia (Iba1), the glioma-bearing mice were perfused with 0.01 M PBS and 4% paraformaldehyde (PFA). Then the brains were collected and fixed in 4% PFA for 12 h, followed by dehydration in 30% sucrose solution for two days. Brain coronal sections with a thickness of 30 µm were sliced using a cryostat (ThermoFisher) and attached to the slides. Brain sections were washed in PBS for 3 times to remove the cryoprotectant solution completely. After blocking (20% normal goat or donkey serum with 0.1% Triton X-100 in PBS) for 1 hour at room temperature, brain sections were incubated with primary antibodies (diluted ratio 1:500): rabbit anti-NeuN (Cat. # AB104225, Abcam), rabbit anti-GFAP (Cat. # RB087A0, Fisher Scientific) and goat anti-Iba1 (Cat. # AB5076, Abcam) for two nights at 4 °C. After another wash by PBS for three times, secondary antibodies (diluted ratio 1:500; Fisher Scientific: donkey anti-rabbit IgG Alexa Fluor 488, Cat. # A21206; donkey anti-goat IgG Alexa Fluor 488, Cat. # A11055) and Hoechst 33342 solution (diluted ratio 1: 2000) were applied for 2 h at room temperature. After washing, all the treated sections were mounted with Fluoromount-G (SouthernBiotech) and imaged by a slide scanner (VS120, Olympus) and spinning disk microscopy (SD-OSR, Olympus).

To stain the pre-and post-synapse, the similar procedure was conducted with the following primary antibodies: Guinea pig anti-VGluT2 (vesicular glutamate transporter 2, Invitrogen, Cat. #OSV00108W) and rabbit anti-PSD-95 (post synaptic density 95 kDa, Invitrogen, Cat. #51-6900), respectively. The following second antibodies were used accordingly: Goat anti-Rabbit IgG FITC (Invitrogen, Cat. #OSV00108W) and goat anti-guinea pig IgG Alexa Fluor 647 (Invitrogen, Cat. #A-21450).

For immunostaining of extracellular matrix (ECM) components, brain sections were incubated overnight at 4 °C with primary antibodies: biotinylated hyaluronan binding protein (HABP) antibody (Amsbio LLC, Cat. #AMS.HKDBC41), collagen IV antibody (Invitrogen, Cat. #PA1-85320), tenascin R (TN-R) antibody (Invitrogen, Cat. #PA5-50580), and anti-Fibronectin antibody (Abcam, Cat. #ab2413). After wasing three times with PBS, brain sections were then inbubated the corresponding secondary antibodies for 2 hours at room temperature: Cy5-Streptavidin (Invitrogen, Cat. #SA1011) targeted at the HABP antibody, and anti-Rabbit IgG antibody conjugated with Alexa Fluor™ 647, specific for the Collagen IV, TN-R, and anti-Fibronectin antibodies. After washing, all the treated sections were mounted with Fluoromount-G.

**Analysis of diffusion coefficient**

Diffusion coefficients were estimated by employing a point-source paradigm with a two-step fitting procedure as described for integrative optical imaging analysis.^4, 5^ First, the fluorescence intensity of each image was fit to a 2-d Gaussian function derived from a model of normal diffusion from an instantaneous point-source.

$I=A exp(-\frac{r^{2}}{\gamma^{2}})$ ,	Equation 1

where $r^{2}=\left( x-x_{0} \right)^{2}+\left( y-y_{0} \right)^{2}$ is the distance from the point-source, (x_0_, y_0_), typically taken as the center of the tornado scan area during photo-stimulation. A and γ are free fitting parameters that were determined by optimizing the fit of Eq. 1 to the fluorescence intensity distribution in an image with the Nelder-Mead algorithm. Second, the fitting parameter γ at each timepoint was taken as a function of time and linear regression was used to estimate the diffusion coefficient from the slope of the line given by Eq. 2.

$\gamma^{2}=4D(t+t_{0})$	Equation 2

The analysis procedure was implemented as a Python program (https://github.com/NTBEL/diffusion-fit) using various numerical and scientific Python libraries, including scikit-image,^6^ SciPy,^7^ NumPy,^8^ pandas,^9^ and Numba.^10^

λ is the tortuosity of the brain extracellular space and is defined by:

$\lambda= \sqrt{D/{D^{*}}}$	Equation 3

where *D* and *D^*^* are the free diffusion coefficient in diluted agarose gel and effective diffusion coefficient *in vivo* or acute brain slices.

**Statistical analysis**

All data were collected in more than triplicate and reported as mean and standard deviation. Two-sample Student’s t-test or one-way ANOVA with Tukey test in Origin 2021 software were conducted to determine p values and statistical significance. Single asterisk (*) indicates p < 0.05, double asterisks (**) indicate p < 0.01, and triple asterisks (***) indicate p < 0.001 as a measure of significance. The Spearman rank correlation ananlysis was performed using Origin 2021 software.

**Code availability**

The custom MATLAB codes for molecular diffusion and data analysis used in this work are available upon request to Z.Q.

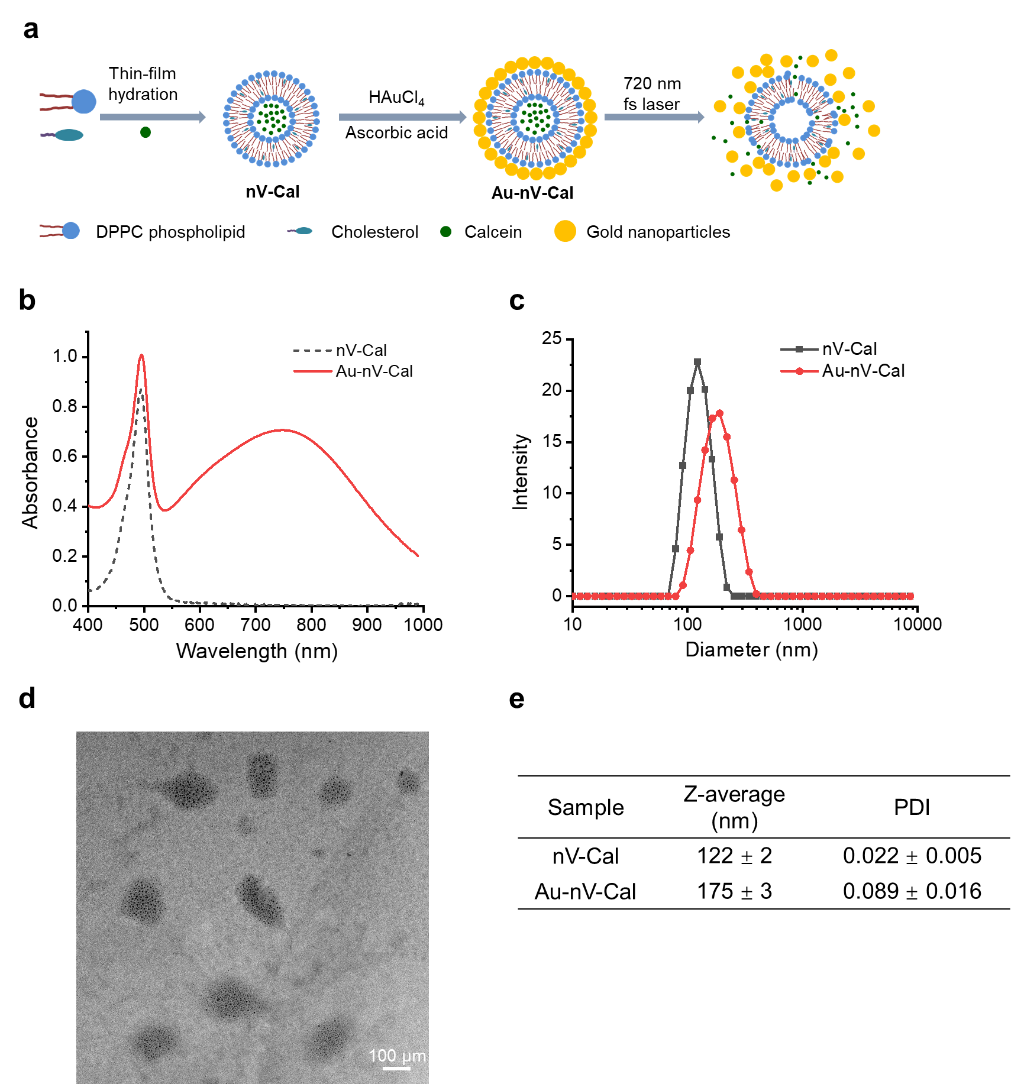

**Figure S1**. Characterization of gold-coated calcein-encapsulated nanovesicles (Au-nV-Cal). (a) Schematic of Au-nV-Cal preparation. (b) UV-Vis spectra of calcein-encapsulated nanovesicles (nV-Cal) and Au-nV-Cal. (c) Size distribution of nV-Cal and Au-nV-Cal measured by dynamic light scattering method. (d) Transmission electron microscope images of Au-nV-Cal. Scale bar: 100 µm. (e) Summary of the particle size and PDI of nV-Cal and Au-nV-Cal.

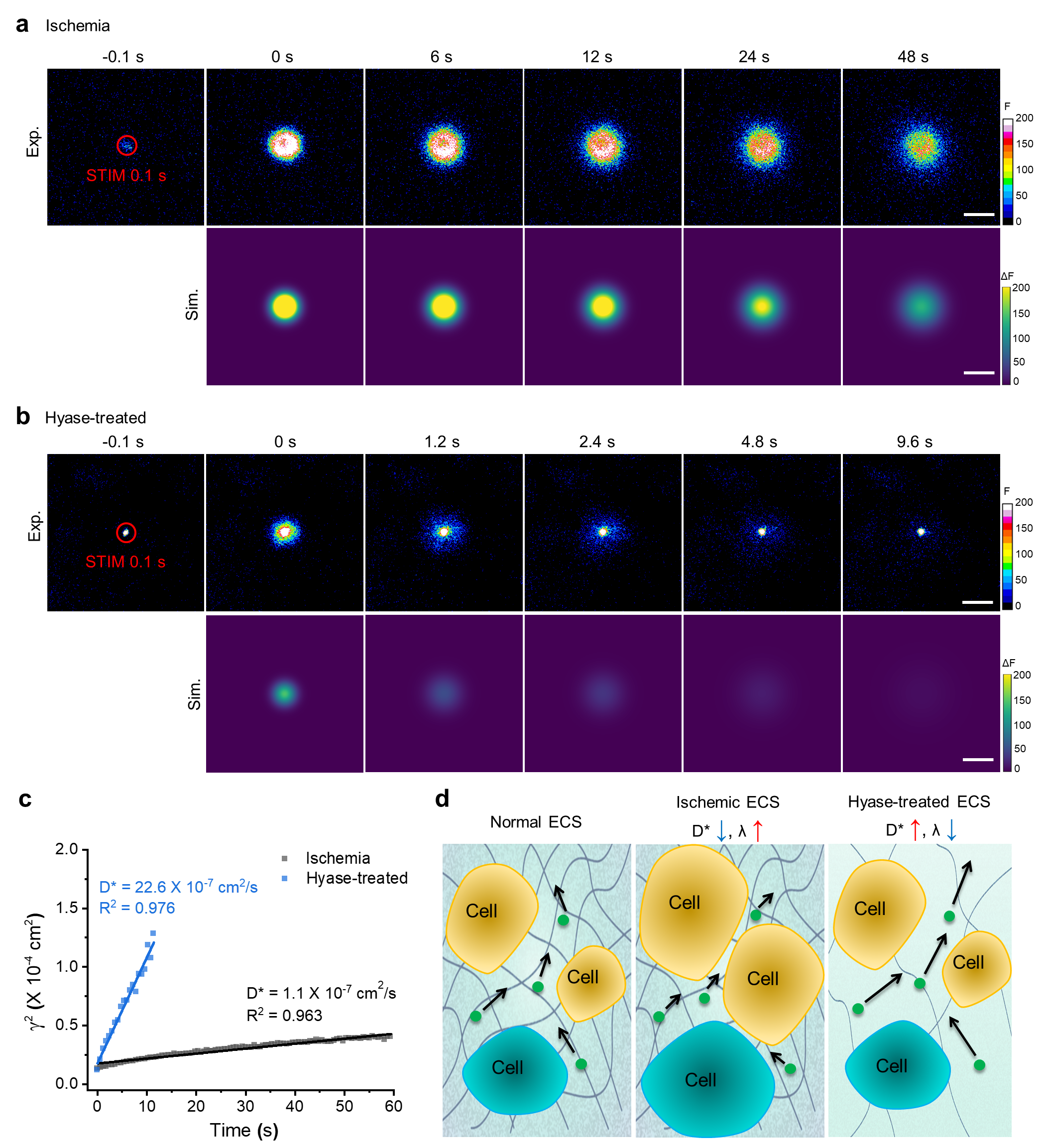

**Figure S2.** Calcein diffusion measurement in the somatosensory cortex of ischemic and hyase-treated mouse brains. (a, b) Two-photon fluorescent images (upper panel) and 2D Gaussian fitted images (lower panel) of calcein diffusion in (a) ischemic and (b) hyase-treated mouse brains. 60-µm-diameter tornado scans (red circle) were performed on Au-nV-Cal for 0.1 s before the diffusion recording. Scale bar: 100 µm. (c) Linear fitting of *γ^2^* versus *t* to get the diffusion coefficient. (d) Schematic of ECS change under different treatments.

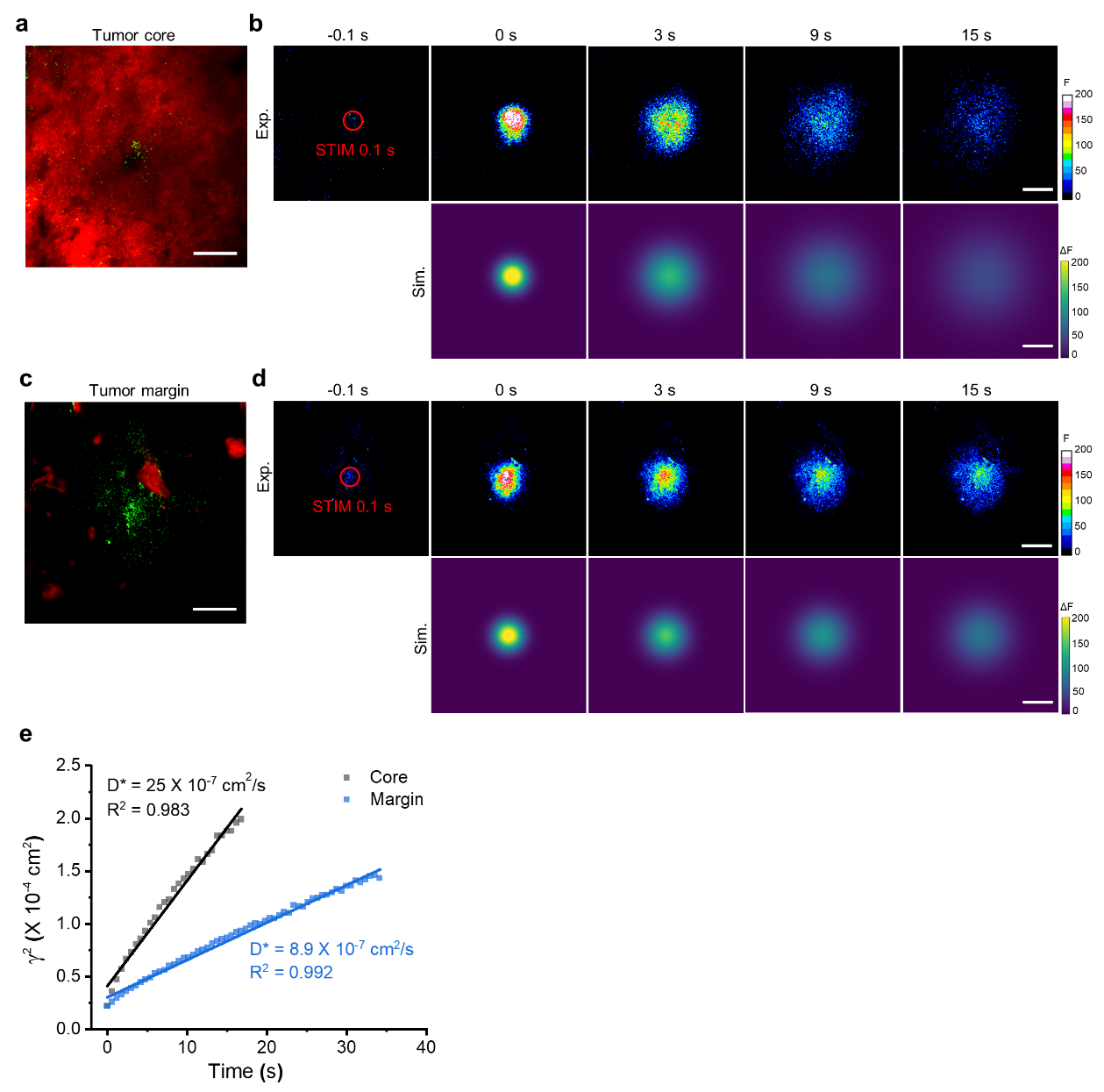

**Figure S3.** Calcein diffusion in 73C GBM. (a,c) Fluorescent image of the core and margin of tdTomato-labelled (red) 73C GBM by two-photon microscopy, respectively. The green indicates the distribution of Au-nV-Cal. Scale bar: 100 µm. (b,d) Two-photon fluorescent images (upper panel) and 2D Gaussian fitted images (lower panel) of calcein diffusion in 73C GBM core and margin, respectively. 60-µm-diameter tornado scans (red circle) were performed on Au-nV-Cal for 0.1 s before the diffusion recording. Scale bar: 100 µm. (e) Linear fitting of *γ^2^* versus *t* to get the diffusion coefficient.

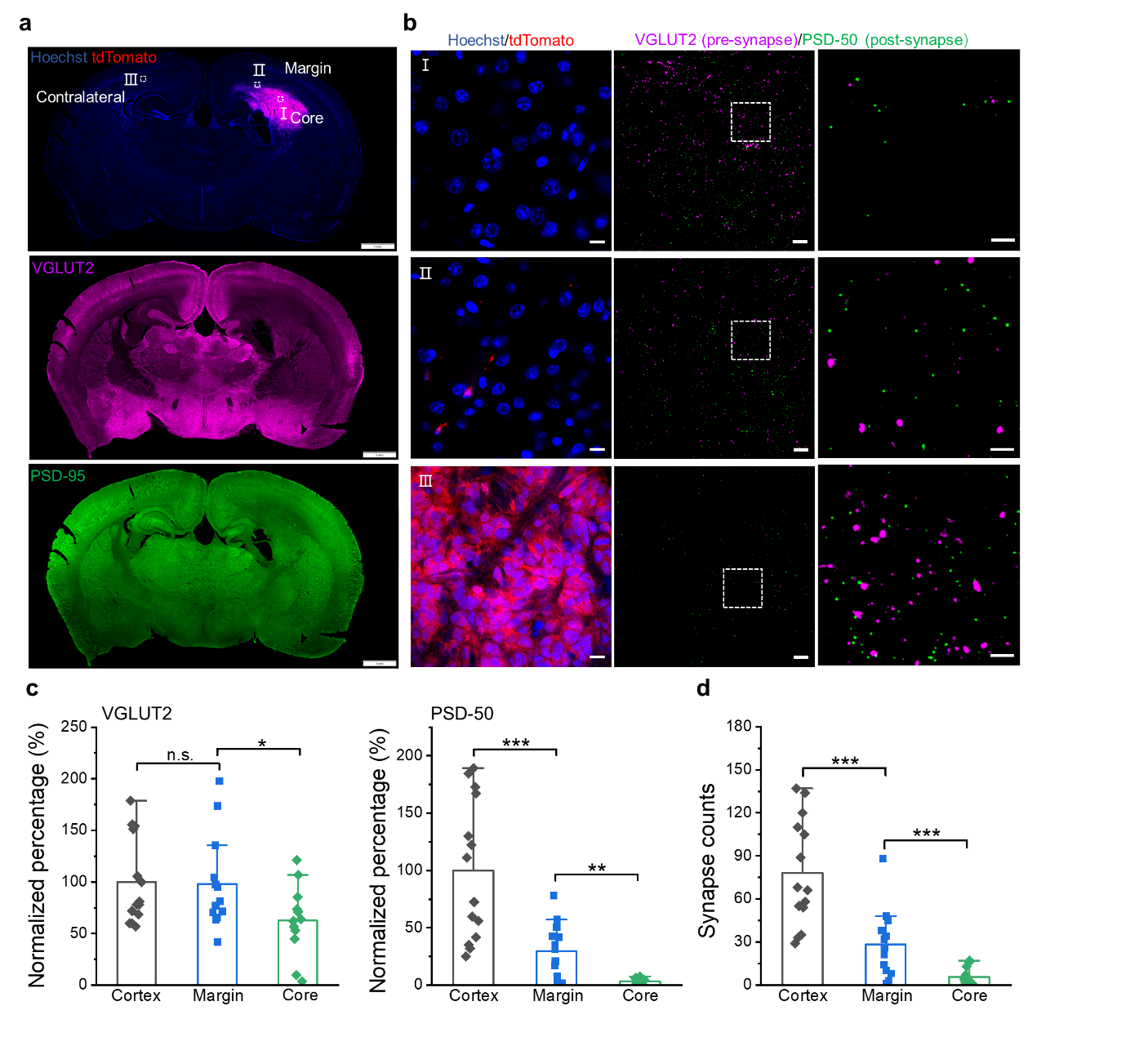

**Figure S4.** Immunostaining of synapses in PS5A1 GBM. (a) Fluorescent images of the whole brain slice immunostained by VGLUT2 (pre-synapse, purple) and PSD-50 (post-synapse, green) in PS5A1 GBM. Glioma cells were labeled by tdTomato (red). Cell nuclei were labeled by Hoechst 33342 (blue). Scale bar: 1 mm. (b) Left two columns： the fluorescent images with high magnification in the three regions labeled in (a); scale bar: 10 μm. Right column: the zoomed-in images; scale bar: 3 μm. (c) The quantitative analysis of the pre- and post-synapse signal in the field of view (133 µm × 133 µm) normalized by the contralateral cortex. (d) The synapsed counts in different regions of PS5A1 GBM. Statistical analysis was performed by two-sample Student’s t-tests. Data are expressed as Mean ± S.D.; ***p* < 0.01; ****p* < 0.001; n.s., not significantly different.

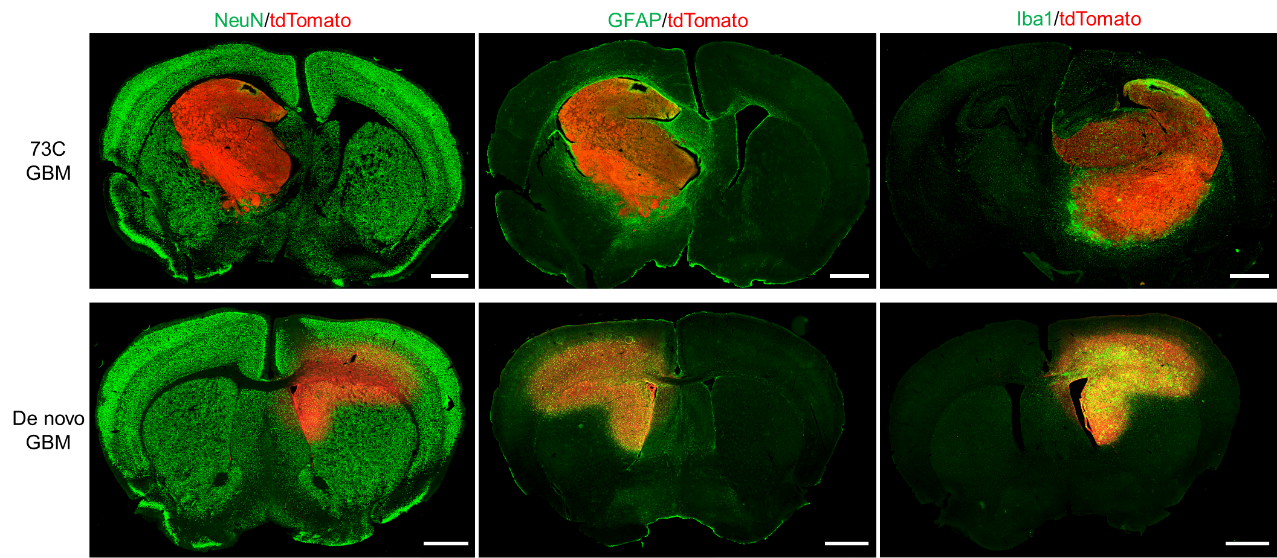

**Figure S5.** Images of immunostained brain cells in the whole slice of 73C and de novo GBMs. Glioma cells were labeled by tdTomato (red). Neurons, astrocytes, and microglia were labeled by Alexa Fluor 488 (green).

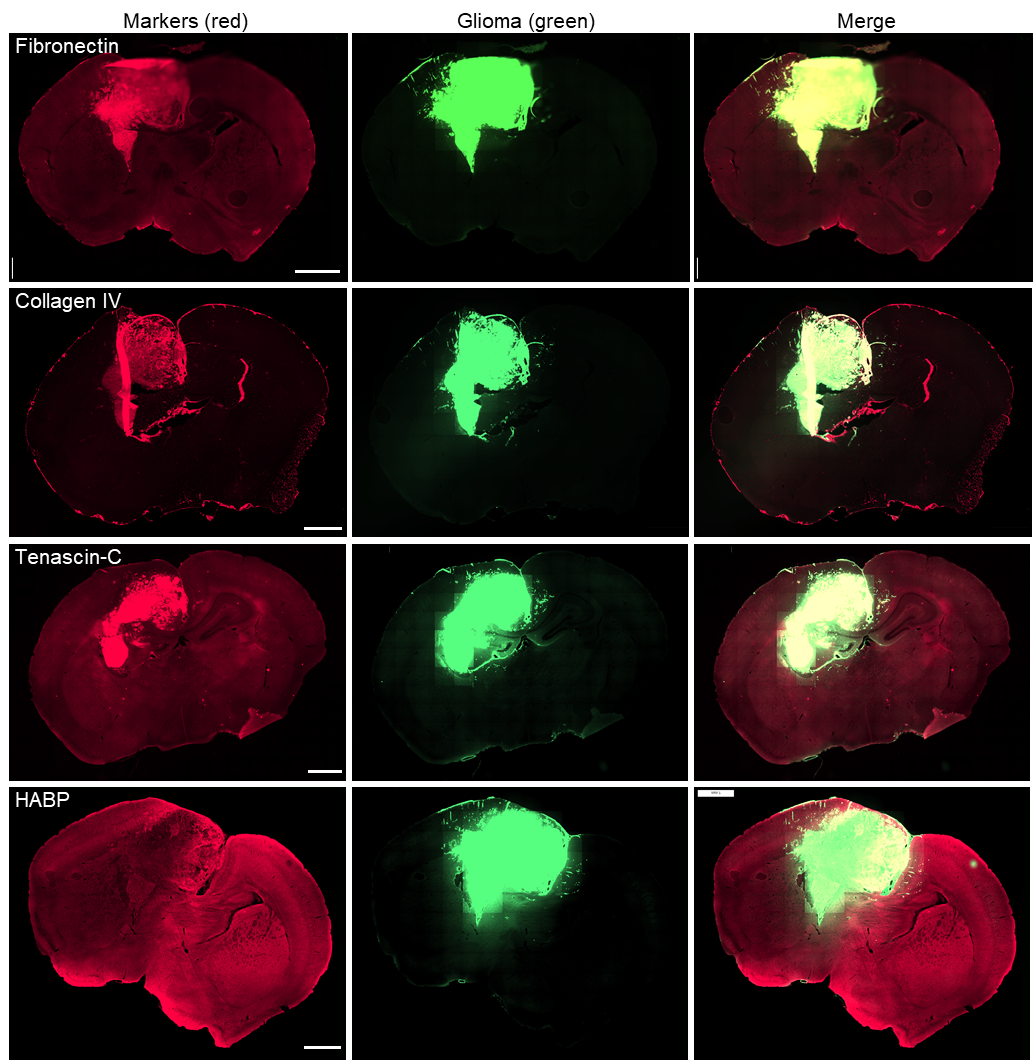

**Figure S6.** Differential expression of extracellular Matrix (ECM) proteins in the 73C GBM. Large field-of-view immunohistochemical images reveal distinct expression of glioma ECM components. Fibronectin, Collagen IV, and Tenascin-C exhibit increased expression in the glioma core relative to the margin, whereas HABP expression remains consistent across both regions. Scale bar: 1 mm.

**Table S1.** Summary of cellular heterogeneity and tortuosity

| Regions& models | Relative number of neurons | Relative number of astrocytes | Relative number of microglia | λ |
| --- | --- | --- | --- | --- |
| Normal cortex | 1.00 | 1.00 | 1.00 | 1.90 |
| PS5A1 margin | 1.00 | 5.69 | 3.88 | 2.30 |
| PS5A1 core | 0.00 | 4.21 | 105.09 | 1.79 |
| 73C margin | 0.85 | 6.01 | 30.45 | 1.89 |
| 73C core | 0.00 | 0.08 | 2.35 | 1.33 |
| de novo margin | 1.08 | 12.51 | 26.38 | 2.59 |
| de novo core | 0.08 | 107.15 | 212.90 | 1.47 |
